# Blocking mechanical airway epithelia damage during asthma progression reverses airway remodelling and hyperresponsiveness

**DOI:** 10.1101/2025.11.10.687569

**Authors:** Dustin C. Bagley, Tobias Russell, Rocio T. Martinez-Nunez, Stefania Marcotti, Jody Rosenblatt

**Author notes:** These authors contributed equally to this work (Dustin C. Bagley, Tobias Russell).

## Abstract

Airway smooth muscle (ASM) hyperresponsiveness is the defining feature of asthma, yet neither its origins nor the processes that establish and maintain it are understood. Here, using a mouse model, we show that hyperresponsiveness arises from ASM realignment into dense, parallel bundles reinforced by excess extracellular matrix, a fundamental remodelling rather than mere amplification of existing contractile ASM. This remodelling is initiated and sustained by a mechanical feedforward cycle. Allergen treatment initiates a contraction-relaxation cycle that realigns ASM into a bronchoconstrictive state. As bronchoconstriction triggers crowding and excess airway epithelial cell extrusion ^1^, the wounding then locks ASM into a hypercontractile architecture that drives further constriction. Gadolinium, a mechanosensitive channel blocker that inhibits extrusion, prevents this remodelling and hyperresponsiveness—unlike the corticosteroid budesonide. Critically, in mice with fully established disease, gadolinium reverses all asthmatic features and restores normal airway function, while budesonide alone cannot. Budesonide, however, synergises with gadolinium to normalise protein and gene expression. These findings reframe asthma pathogenesis as a mechanically driven, pharmacologically reversible process — suggesting that asthma may be curable rather than merely treatable.

---

2800 years ago, Homer first mentioned ‘asthma’ as the extreme breathlessness of warriors in battle, with Hippocrates (400 BC) adopting it medically to describe his patients with breathing difficulty ^2,3^. Today, asthma remains a common disease, predicted to now affect more than 400 million people annually, with half a million associated deaths. The bronchoconstriction of an attack causes not only breathlessness, but also increased mucus, airway epithelial damage, and immune infiltration ^1^. Current frontline therapies treat the immune response with inhaled corticosteroids that target airway inflammation and **ß**2 adrenergic receptor agonists that reverse bronchoconstriction by relaxing airway smooth muscle (ASM) ^4,5^. Severe asthmatics (∼5% of patients) continue to experience asthma exacerbations and require intermittent oral corticosteroids with high dose inhaled corticosteroids to adequately manage their disease. Despite the recent introduction of targeted anti-T2 biologic therapies, clinical remission rates are only in the range of 20-30% highlighting a significant unmet need ^6-8^.

While new therapeutics improve asthma symptom treatment and slow disease progression, none modify disease outcomes over time or cure it ^1,9^. It is important to note that most asthma therapies control immune cell response, rather than preventing the main unifying feature – bronchoconstriction. Yet, controlling the propensity to bronchoconstrict is the most critical issue, as we found that the mechanics of bronchoconstriction causes airway wounding and inflammation due to too excessive epithelial cell extrusion and mucus release ^1^. A key unanswered question is why only asthma patients hyperconstrict in response to asthma triggers. While presumably ASM amplification can promote hyperresponsiveness, we lack a biophysical picture of what causes the changes that cause hyperconstriction in the first place. Without understanding how and why airways become hyperresponsive, our ability to regenerate healthy, non-responsive airways remains limited.

## House dust mite, but not methacholine, remodels airway smooth muscle in mice

To investigate ASM remodelling in our asthma model, we compared airways from control mice versus mice immune-primed with house dust mite (HDM) five times per week for three weeks using a standard protocol (fig. S1A) that routinely produces type 2 inflammation and airway hyperresponsiveness ^1,10^. By filming ex vivo lung slices treated with methacholine (MCH) to induce bronchoconstriction, we confirmed that HDM-primed slices hyper-constricted, causing pronounced epithelial extrusion, buckling, and airway lumen occlusion, as seen previously ^1^, whereas unprimed airways had little to no response (SFig. 2A-C, movies S1 and S2).

To investigate how ASM changes in response to HDM-priming, we stained ex vivo lung slices for actin and imaged longitudinal sections by confocal microscopy (Fig. 1A). We found that healthy, unprimed lung slices formed sparse, lattice-like ASM bundles, whereas those from HDM-treated mice became tightly aligned with short gaps between actin bundles. Quantification of the average distance between actin bundles for each airway confirmed that HDM-treated airways have significantly shorter spaces between ASM bundles compared to controls (Fig. 1B). Order parameter measurements (the degree of orientational alignment; see methods) indicated that airway smooth muscle became more aligned and ordered into parallel bundles in HDM-treated lungs, compared to healthy controls (Fig. 1C). Interestingly, the parallel bundling also coincided with less cross-hatching of each bundle, as exemplified by red lines on ASM bundles that intersect with each other (Fig. 1D). Histological analysis also revealed that the combined ASM and basement membrane thickens in HDM-treated lungs, compared to controls (Fig. 1E and F), seen in human asthma pathological samples ^11^. Notably, mathematical modelling has linked airway smooth muscle orientation and submucosal thickening with bronchoconstriction ^12,13^. Thus, we found a strong correlation with ASM realignment into tightly-space parallel bundles, which may enable ASM bundles to constrict together without restraint, compared to healthy airways where smooth muscle contraction may be impeded by contracting against each other in lattice-like configurations.

**Figure 1.**
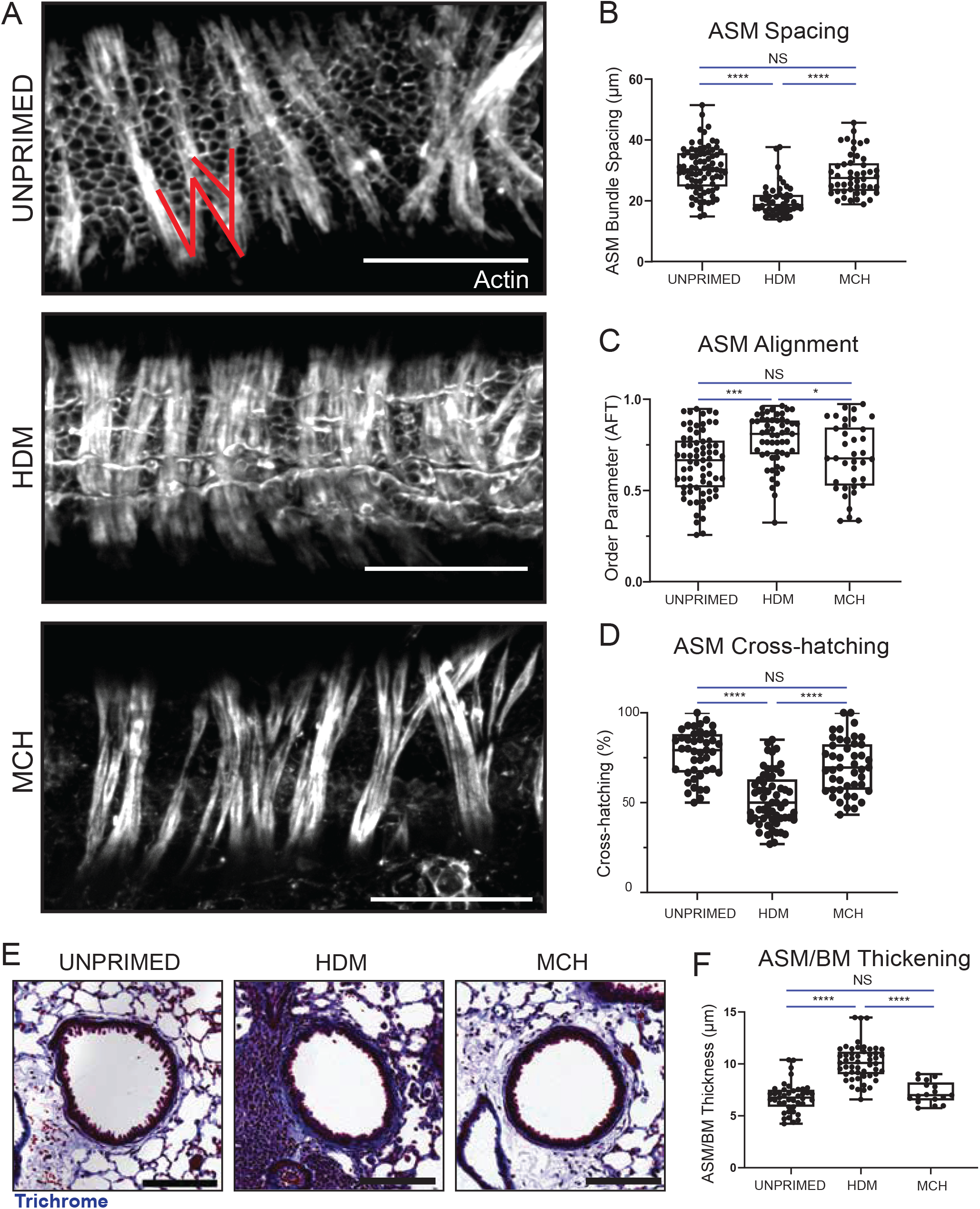
Airway smooth muscle remodeling in hyperresponsive airways. (A) Confocal projections of phalloidin-stained lung slices highlighting ASM amplification and alignment in HDM treated mice, healthy controls, and MCH primed mice (Scale bars, 100 um). Quantification of ASM spacing (B), alignment (C) and percentage of cross-hatching (red lines on (A)) (D) from confocal projections of phalloidin-stained lung slices represented in (A) unprimed: 71 airways from n=4; HDM: 55 airways from n=5; MCH: 37 airways from n=3 (*p<0.01; ***p<0.0005; ****p<0.0001 from a Mann-Whitney test). (E) Representative airways from control, HDM-, and MCH-primed mice stained with mason’s trichrome, highlighting ASM and basement membrane in blue (Scale bars, 100 um). (F) ASM and BM thickness measurements using trichrome-stained histology images (E) showing a significant increase in HDM primed mice, but not in MCH primed lungs, compared to controls (****p<0.0001 from a Mann-Whitney test).

While HDM-priming is known to produce hyperresponsiveness, it is not clear how it does so and whether simply repeatedly inducing ASM contraction alone might lead to a similar response. Numerous studies have demonstrated that mimicking the mechanics of a hyperresponsive event in vitro can cause and reinforce asthma signatures ^14-16^. Therefore, we next tested if experimentally activating ASM contraction with MCH alone causes the same ASM realignment from HDM-treatment using a protocol that uses a ramp-up MCH challenge instead of HDM-treatment (fig. S1B&C). MCH priming neither caused MCH-induced hyperconstriction in ex vivo lung slices nor did it impact ASM structure, remaining helical and sparsely spaced, with similar cross-hatching to unprimed mice (Fig. 1A-F). Further, histology revealed no immune infiltration or change to ASM thickness (Fig. 1E and F).

### HDM realigns airway smooth muscle through mechanically-induced signalling

We next investigated why HDM-, but not MCH-priming, drives airway remodelling. While the immune response to HDM is well documented, little is known about how it mechanically impacts healthy airways. Thus, we first imaged how lung slices from unprimed mice respond to the first dose of HDM. Remarkably, we found HDM caused a strong mechanical response in more than half of all unprimed airways filmed, regardless of their size. Within minutes of the first HDM dose, we noted rapid cycles of constriction and relaxation lasting for >30 minutes that could reduce lumen areas as much as 50% (Fig. 2A and B, movies S3 and S4). The mechanical crowding with HDM, however, did not cause noticeable rates of extrusion, possibly from its transient or cyclical nature. Importantly, cyclic stretch is a known driver of ASM alignment and proliferation that could potentially stretch and pull smooth muscle bundles into tighter alignment, creating a pathological architecture that allows for hyper-constriction ^17,18^. By contrast, MCH had little effect on airway lumen areas over time when given to healthy airways (Fig 2A and B, movie S5). As a hyper-response to a MCH challenge is the diagnostic test for asthma ^19^, this was not surprising. Thus, HDM, but not MCH, causes a robust mechanical response of unprimed healthy airways that could help realign ASM into its pathological configuration.

**Figure 2.**
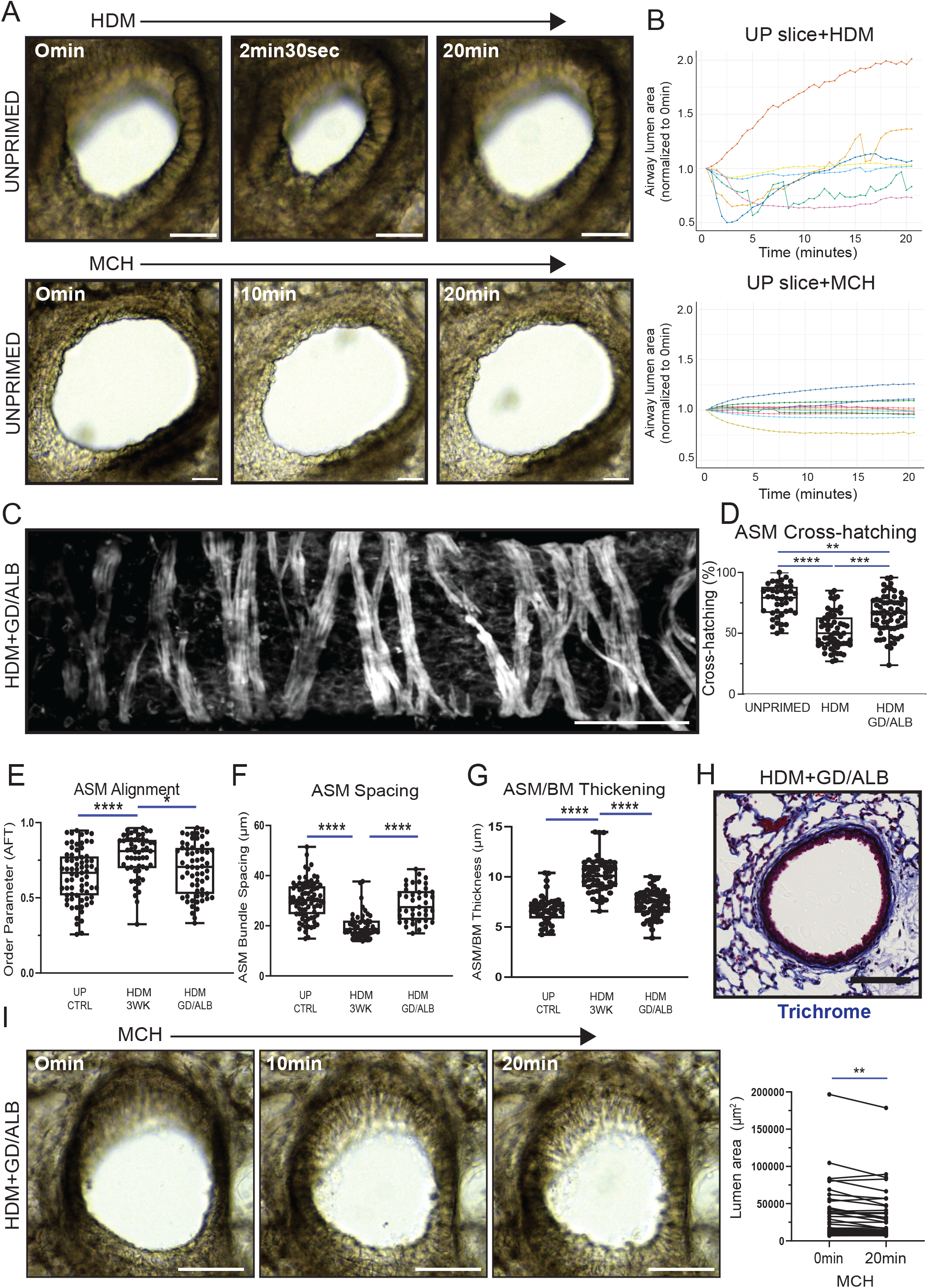
Gadolinium treatment with onset of HDM priming prevents ASM remodelling. (A) Movie stills of 1µg/µL HDM (top) or 500 mg/mL MCH (bottom) responses in healthy, unprimed ex vivo lung slices over time (Scale bars, 50 µm). (B) Quantification of airway lumen areas every 30 secs for 20 mins from unprimed lung slices treated with either HDM (n=7 airways) or MCH (n=8 airways), normalizing lumen area to 1.0 at time 0 min. (C) Representative confocal longitudinal projection of an airway stained for actin to highlight ASM from mice primed with HDM+GD/ALB (Scale bars, 100 µm). Quantification of the percentage of ASM cross-hatching (D), alignment (E), spacing (F) and ASM/BM thickness (G) from mice primed with HDM, HDM+GD/ALB and healthy, unprimed con trols (*p<0.05, ****p<0.0001 from a Kruskal-Wallis test). (H) Representative airway from GDM+GD/ALB mice stained with mason’s trichrome. (I) Movie stills from ex vivo lung slices from HDM+GD/ALB primed mice treated with 500 mg/mL MCH (Scale bars, 50 µm), where constriction is quantified from measuring the lumen area at time 0 min and 20 mins following 500 mg/mL MCH treatment (45 airways from n=4, **p<0.001 from a Wilcoxon pairs-matched signed rank test).

Yet, since one treatment of HDM is not sufficient to induce hyperresponsive airways, we next tested if continued damage from the many HDM exposures needed to prime airways may fix them into a hyperresponsive configuration. Based on our previous findings that hyper-constriction causes excess extrusion, wounding the airways ^1^, one possibility is that damage could reconfigure ASM into a chronically contracted state to reduce the surface area that would need to be covered by fewer epithelial cells. Thus, we tested if blocking epithelial cell extrusion with a potent, generic stretch-activated channel (SAC) inhibitor, gadolinium (GD) (10), during HDM priming could prevent airway remodelling and inflammation. We first tested the safety of administering GD on a continuous basis during HDM priming. To assay if Gd^3+^Cl_3_ (non-chelated) accumulates in the lung or other tissues after multiple administrations, we intranasally instilled 10μM Gd^3+^Cl_3_ once a day for five days (fig. S1E), sacrificing mice at 30 min or 24 hrs after the last instillation and harvesting lungs (with trachea), liver, kidney, and part of the gastrointestinal tract (GI) (fig. S3A). Using inductively coupled plasma mass spectrometry (ICP-MS) to precisely measure total Gd^3+^ concentrations, we found that less than 0.6 µg/g Gd^3+^ (sub parts per million, <10% of the total dose over 5 days) was within the lungs 30 min after the last Gd^3+^Cl_3_ administration, which decreased further after 24 hrs (fig. S3A and B). Negligible Gd^3+^ was detected in the liver, kidney, and GI. Expectedly, no Gd^3+^ was detected in the lungs of untreated control mice. Thus, intranasal instillation of uncaged Gd^3+^Cl_3_ does not accumulate appreciably in the lungs or other tissues.

Having established that repeated doses do not accumulate or cause appreciable harm to mice, we primed mice for three weeks with HDM ±GD and albuterol (ALB), a short-acting **β**2 adrenergic receptor agonists that relaxes ASM but does not protect against epithelial damage or inflammation (fig. S1D) ^1,20^. Importantly, we found that adding GD/ALB during HDM-priming prevented ASM remodelling, preserving the smooth muscle and matrix architectures seen in unprimed airways (Fig. 2 C-I). Accordingly, preserving the ASM architecture reduces airway response to MCH, compared to HDM-priming alone (Fig. 2H, movie S6), further confirming the link between the ASM remodelling and airway hyperresponsiveness. Moreover, as expected from our previous studies ^1^, the reduced constriction in response to GD/ALB/HDM priming resulted in significantly reduced airway epithelial extrusions and denuding following an ex vivo MCH challenge (fig. S2C-E). Along with preserving the ASM and matrix architecture, we found that HDM+GD/ALB prevented the loss of ciliated (FoxJ1-positive) cells (fig. S2F and G) and immune cell infiltration (measured by H&E staining) that occurs in HDM-primed alone (fig. S2H and I). Together, our data indicates that preventing epithelial damage with gadolinium during HDM priming prevents airway remodelling and hyperresponsiveness, suggesting a link between wound healing, remodelling and hyperreactive airways.

### Blocking ongoing epithelial damage and inflammation synergistically reverses airway hyperresponsiveness

Since asthma treatments only begin once an individual receives a diagnosis, we investigated if gadolinium could reverse airway remodelling and asthma features after they begin. We compared our GD treatments to budesonide (BUD), a corticosteroid anti-inflammatory – the cornerstone ‘preventer’ long-term therapy used to reduce asthma attacks and their severity. Given that three weeks of HDM-priming caused ASM structural changes, MCH hyperresponsiveness, and epithelial damage (Figs. 1 and 2), we first primed mice with HDM for three weeks and then intranasally instilled mice with GD, BUD, or both for another two weeks along with HDM (fig. S1F). We continued HDM exposure during treatment, since asthmatics are likely to encounter allergen triggers throughout any real-world intervention period.

Strikingly, movies from ex vivo methacholine challenges revealed that both the HDM-alone group and the HDM+BUD group were highly hyperresponsive, exhibiting excessive epithelial extrusion and damage. In stark contrast, adding GD during the final two weeks of HDM priming dramatically reduced hyperresponsiveness, restoring them to levels seen in unprimed mice (movies S7-10). Quantifying changes in airway lumen area before and after challenge, showed that two weeks of GD treatment during ongoing HDM priming (± BUD) greatly reduces bronchoconstriction in most airways (Fig. 3A). By contrast, the additional two weeks of HDM priming in HDM controls slightly increases hyperresponsiveness (Fig. 3A). In agreement with our previous study, the reduction in airway hyperresponsiveness corresponds with a marked decrease in epithelial extrusion and denudation, reaching levels seen in unprimed mice (Fig. 3B–D). Remarkably, BUD alone had no effect on any of these metrics (Fig. 3B–D), despite being considered a preventative agent.

**Figure 3.**
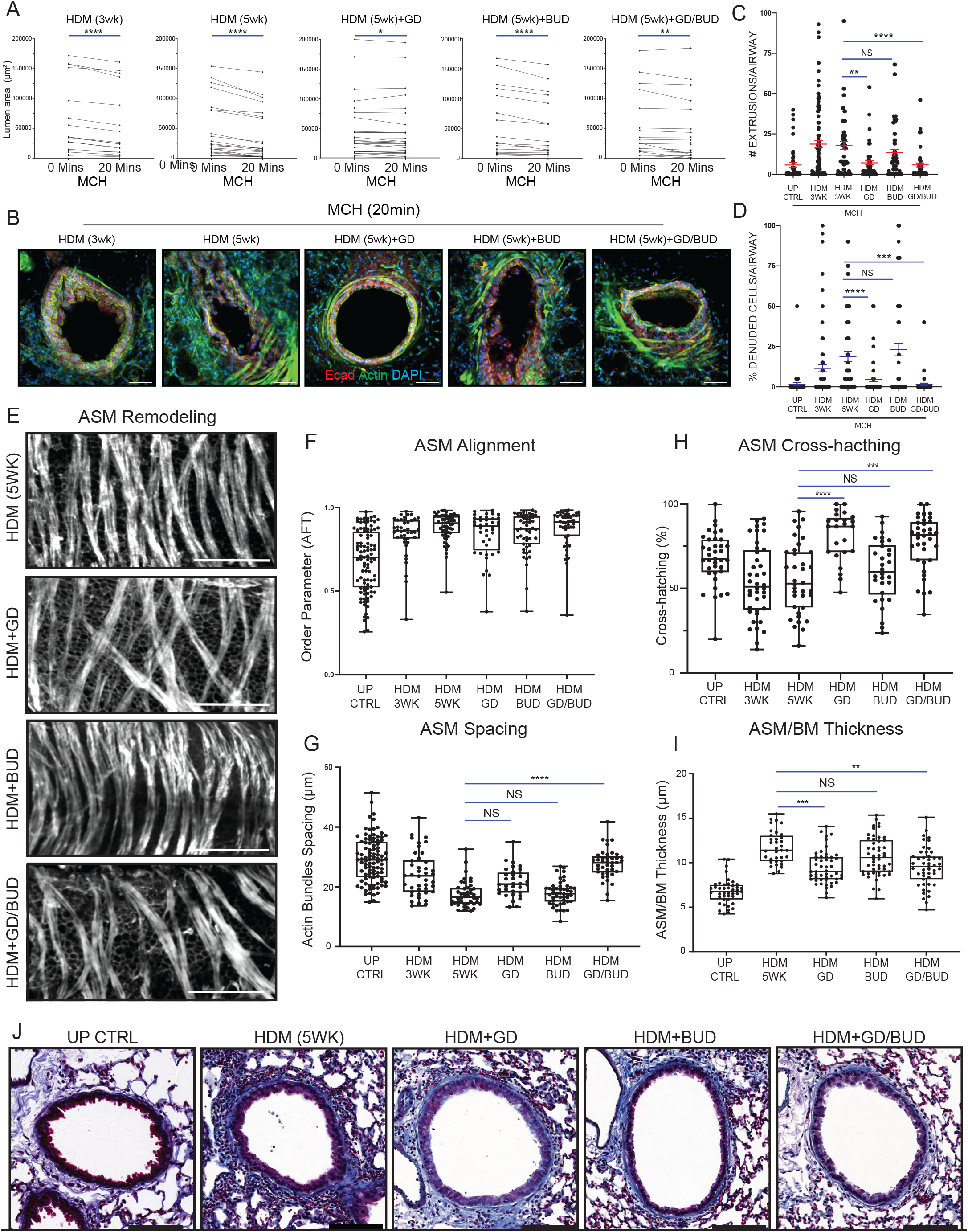
Gadolinium reverses airway remodelling and hyperresponsiveness. (A) Representative confocal projections of airways stained for actin, showing ASM remodelling and alignment with quantification of ASM alignment (B), augmentation (C), percentage of cross-hatching (D), and ASM/BM thickness (E) from 6 unprimed mice (95 airways); 6 3wk HDM mice (42 airways); 6 5wk-HDM mice (44 airways); 6 HDM+GD mice (34 airways); 6 HDM+BUD mice (48 airways); and 5 HDM+GD/BUD mice (41 airways), with *p<0.05, **p<0.001, ***p<0.0005, ****p<0.0001 from a Kruskal-Wallis test. (F) Trichrome-stained histological sections from mice primed with HDM -/+ treatments. (G) Measurements of ex vivo lung slice bronchoconstriction following 500 mg/mL MCH from mice primed with HDM (3wks) (24 airways from n=3), HDM (5wks) (26 airways from n=3), HDM+GD (29 airways from n=3), HDM+BUD (20 airways from n=3), or HDM+GD/BUD (23 airways from n=2) by measuring lumen area at 0 min and after 20 min. (*p<0.05, **p<0.001, ****p<0.0001, from a Wilcoxon pairs-matched signed rank test). (H) Confocal projections of representative airways following an ex vivo MCH challenge immune-stained for the epithelium (Ecad), actin (phalloidin), and DNA (DAPI) (Scale bars, 50 µm), quantified for epithelial cell extrusion (I) and percent denuding (J) per airway (unprimed: 47 airways from n=3; HDM (3wks): 118 airways from n=9; HDM (5wks): 73 airways from n=6; HDM+GD: 76 airways from n=6; HDM+BUD: 73 airways from n=6, HDM+GD/BUD: 71 airways from n=5) *p<0.05, **p<0.001, ***p<0.0005, ****p<0.0001 from a Mann-Whitney test.

Importantly, HDM priming with GD reversed the airway remodelling that we defined by smooth muscle bundle thickness, cross-hatching and alignment (order parameters), and BM thickening (Fig. 3 E-J). BUD on its own did not alter any of these measurements compared to HDM priming alone at 5 weeks (Fig. 3 E-J). GD ± BUD also significantly decreased the BM thickness, compared to 5-weeks of HDM priming, whereas BUD on its own had no impact (Fig. 3I and J). Amplification of the underlying matrix may tether airway smooth muscle into a tightly aligned configuration that contributes to hyperresponsiveness. Interestingly, GD also reduces matrix buildup. We, therefore, observe a very strong correlation between airway hyperresponsiveness and the remodeling of airway smooth muscle into tight, parallel-aligned bundles reinforced by matrix, a pattern reversible with GD. This finding suggests that such remodeling results from ongoing mechanical damage, which anti-inflammatory drugs do not alter. Importantly, preventing ongoing mechanical damage and epithelial wounding allows us to restore the airway smooth muscle to a less-responsive state, even after remodeling is established.

### Gadolinium protects against bronchoconstriction-induced inflammation whereas budesonide does not

As inflammation also contributes to other asthma features, we compared how GD versus BUD impacts over-all lung morphology by H&E histology. We found both GD and BUD greatly reduced immune infiltrates and macrophage immunostaining compared to HDM alone, with the combination having the greatest effect (Fig. 4A and B and fig. S4A and B). GD, independently of BUD, could also restore ciliated cells back to control unprimed levels (Fig. 4C and D). Thus, GD/BUD treatment can reverse the damage and inflammation seen in immune-primed mice.

**Figure 4.**
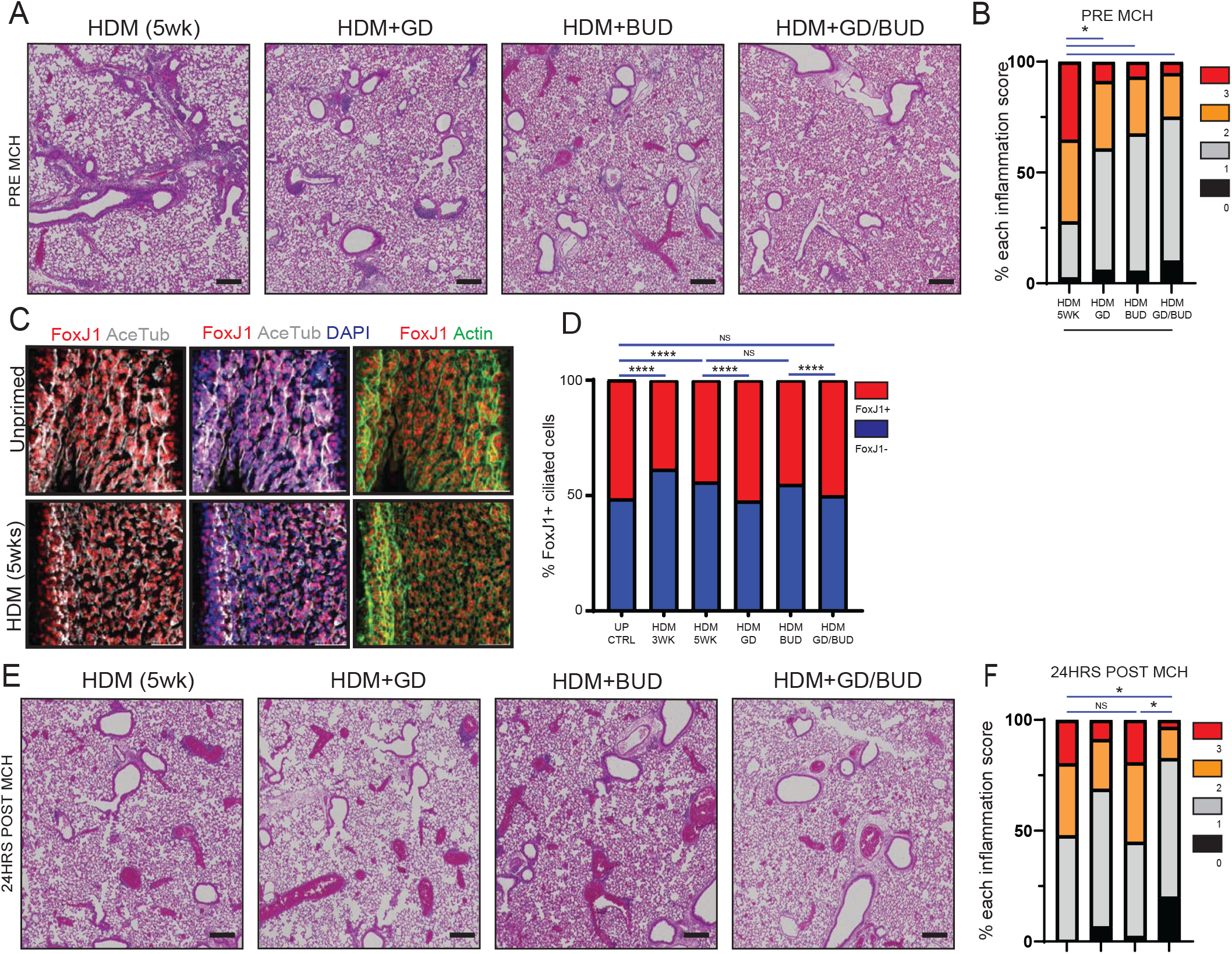
Gadolinium protects against bronchoconstriction-induced inflammation whereas budesonide does not. (A) H&E lung sections from mice treated with HDM for three weeks before priming another two weeks with HDM + GD (n=5), BUD (n=5), GD/BUD (n=5), or control (n=5) that have been quantified for immune infiltration in (B) with black/0 being little-to-no inflammation and red/3 severe inflammation (n=5 per treatment group, *p<0.05 from a Chi-squared test). (C) Representative confocal projections of ex vivo monolayers immune-stained for FoxJ1 (ciliated cell marker) and actin from healthy and HDM (5wks) primed mice (Scale bars, 25 µm). (D) Quantification of the number of ciliated cells across all treatment groups (n=3; ****p<0.0001 from a Chi-squared test). (E) H&E lung sections from mice treated with HDM for three weeks before priming another two weeks with HDM + GD (n=5), BUD (n=5), GD/BUD (n=5), or control (n=5) 24hrs post in vivo MCH challenge that have been quantified for immune infiltration in (F) (n=5 per treatment group, *p<0.05 from a Chi-squared test).

While both GD and BUD protected lungs from basal inflammation during the priming process, the epithelial damage from methacholine challenge seen in HDM ± BUD predicted that only GD might prevent airway inflammation after a MCH challenge. To test this hypothesis, we MCH challenged mice primed with HDM ± GD, BUD, or GD/BUD and analysed lung slices the following day. Although being termed a preventative treatment, continued HDM priming with BUD had no impact on the bronchoconstriction-induced inflammatory response compared to control HDM-primed mice. However, GD-priming (with HDM) alone for two weeks completely blocked this inflammatory response, with GD/BUD decreasing levels further still (Fig. 4E and F). Together, these findings indicate that preventing airway epithelial damage not only reduces basal inflammation but also restores the airway smooth muscle to a non-responsive, unprimed state. As a result, upon challenge, the airways do not bronchoconstrict, thereby avoiding acute epithelial damage and immune cell infiltration, consistent with previous reports ^1^.

### Gadolinium and budesonide synergistically reverse asthmatic lung molecular signatures

Our tissue-based analysis suggested that GD not only reverses airway hyperresponsiveness but also mitigates the overall damage and inflammation caused by repeated allergen exposure. To investigate how GD and BUD affect the protein/gene expression and signaling pathways in airways, we analyzed the transcriptomes and proteomes of whole lungs from healthy controls and mice following three weeks of HDM priming, with or without two additional weeks of BUD and/or GD treatment. RNA-seq cross-group comparisons revealed 1940 differentially expressed genes (p-adjusted < 0.01). We found that GD or BUD individually partially reversed the HDM priming signatures, with the combination nearly completely reversing them from 5 weeks HDM priming (Fig. 5A). By contrast, HDM controls with 2 weeks of additional HDM alone priming only exaggerated the 3 week HDM priming signatures, underscoring the magnitude of GD/BUD reversal (Fig. 5A). Gene ontology analysis revealed that GD/BUD predominantly modified cytokine and chemokine signalling, inflammation, and tissue remodelling pathways (fig. S5A). Importantly, GD/BUD decreases collagen I and V, which have well-documented roles in subepithelial thickening and fibrosis in asthma. While both GD and BUD reduced macrophage- and interleukin-associated genes, consistent with reduced inflammation, GD had a greater impact than BUD (Fig. 5B) ^21^. These findings align with the observed decrease in immune cell infiltration (Fig. 4A and B, fig. S4A and B) and restoration of BM thickness (Fig 3E and F). Interestingly, muscle-related genes were upregulated following GD/BUD treatment. Although many of these genes are typically associated with cardiac muscle, the origin of the smooth muscle involved in airway smooth muscle regeneration back to an unresponsive state remains unclear. Thus, these changes may reflect merely alterations in blood vessels or may suggest that airway smooth muscle regenerates from alternative cell types or through different genetic pathways than previously thought.

**Figure 5.**
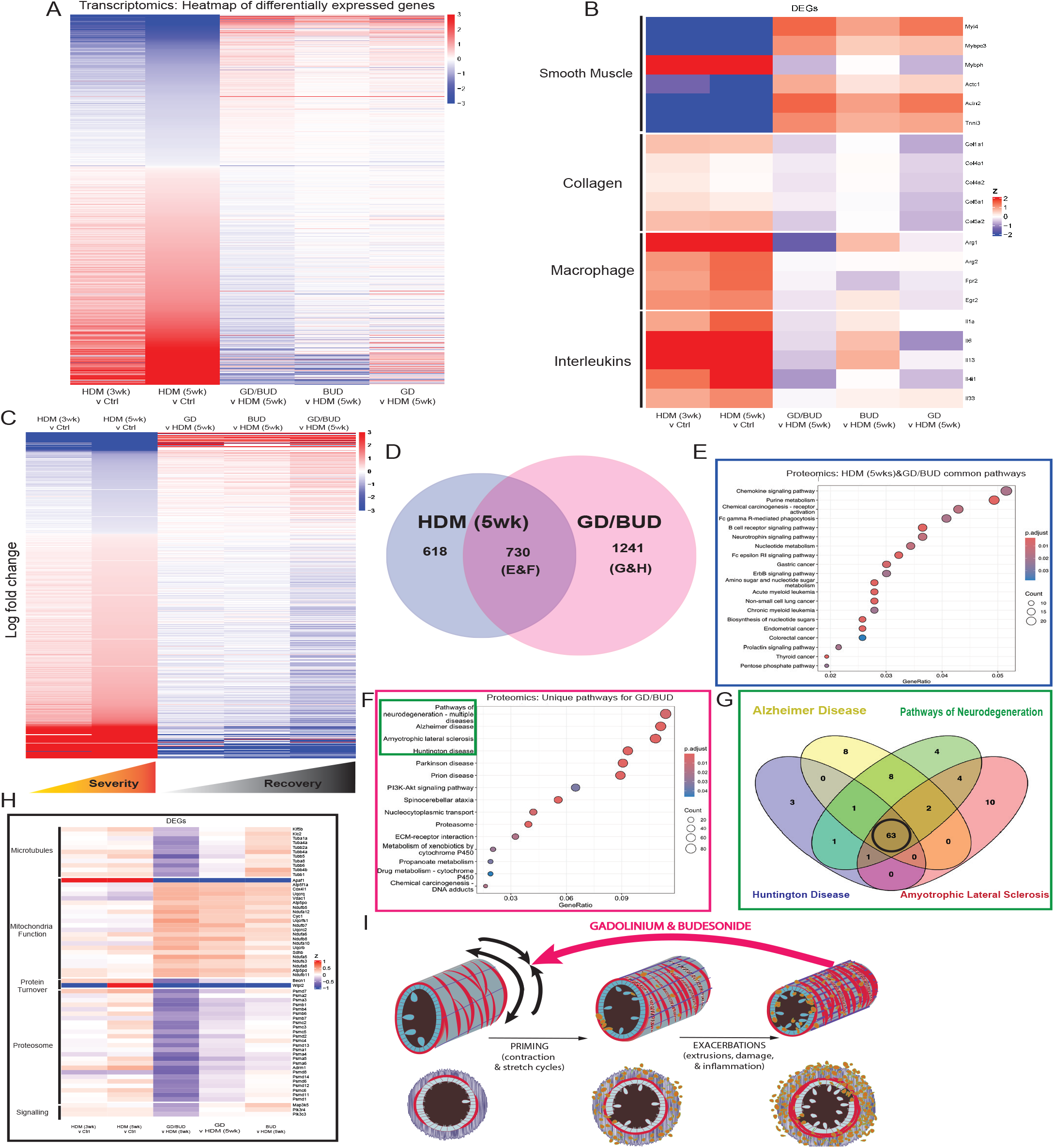
Gadolinium and budesonide synergistically reverse asthmatic molecular signatures. Heatmap of log2 fold differentially expressed genes in HDM (5wk) vs control and treatment groups against HDM (5wk) (both p-adjusted < 0.01). (B) Heatmap of log2 fold changes of candidate differentially expressed genes in HDM (5wk) vs control and treatment groups against HDM (5wk) (p-adjusted < 0.01). (C) Heatmap of log2 fold changes of differentially expressed proteins from 5-week HDM priming vs control unprimed (p-adjusted < 0.05) compared to different treatments. (D) Venn diagram showing proteins in 5-week HDM priming (HDM (5wk) vs control) versus gadolinium + budesonide (GD/BUD vs HDM (5wk)) treatment (p-adjusted < 0.05). (E) Dot plot showing KEGG pathways of differentially expressed proteins from Venn diagram of HDM (5wk) vs control and GD/BUD vs HDM (5wk). (F) Dot plot of KEGG pathways from differentially expressed proteins uniquely modified by gadolinium and budesonide (GD/BUD vs HDM (5wk). GO: Gene Ontology. (G) Venn diagram of differentially expressed proteins (p-adjusted < 0.05) from top four neurodegenerative pathways. (H) Heatmap of log2 fold changes of 63 common proteins from HDM (5wk) vs control (G, black circle, p-adjusted < 0.01). (I) A feed-forward cycle drives airway disease: mechanical stress and injury trigger ASM alignment and amplification, fuelling extrusion and inflammation, which feeds forward further remodelling. GD/BUD synergistically breaks this cycle by blocking epithelial damage and inflammation, restoring lung health.

Proteomics corroborated the patterns observed in the transcriptomic analysis, further highlighting a link between HDM priming and wound healing. Similar to transcriptomics, protein expression changes from GD or BUD alone partially reversed the asthmatic signature from 5-week HDM priming, with the GD/BUD combination producing a more pronounced, synergistic reversal compared to HDM priming alone (Fig. 5C). Using the Venn diagram in Fig. 5D, we found that the protein changes shared between HDM priming and GD/BUD-treated groups were markedly reversed relative to 5-week HDM priming (fig. S5B and Fig5E, blue box). These pathways included immune-related processes (chemokine, B cell, neutrophil, and mast cell signalling) as well as nucleotide metabolism, associated with macrophage activation ^22^.

Pathways uniquely altered by GD/BUD treatment from the Venn diagram (Fig. 5D) are dominated by airway structural pathways (Fig. 5F, pink box), including <u>e</u>xtra<u>c</u>ellular <u>m</u>atrix (ECM)-receptor interactions and proteasomal markers, and an apparent enrichment for neurodegenerative disease pathways (Fig. 5F, green box). However, further investigation of the neurodegenerative proteins (Fig. 5G, black circle) revealed they primarily reflect general features of wound healing, remodelling, and homeostatic cellular functions, rather than any proteins specific to neuronal diseases (Fig. 5H). Specifically, GD/BUD reduced protein turnover and proteosome signatures, as well as microtubule dynamics, motors, and PIK3/MAP3K kinase signalling, which may be reflective of cell and protein degradation and proliferation, commonly occurring during wound healing. By contrast, GD/BUD enhanced mitochondrial function, associated with differentiation and down-regulated in proliferative wound healing states that depend more heavily on glycolysis ^23,24^. Notably, proteasome downregulation has previously been linked with steroid-dependent reduced airway inflammation ^25,26^. Proteomics also revealed that GD/BUD increased ECM-receptor interactions, which could reflect necessary interactions for regenerating the ASM and matrix, likely together creating the correct architectures for non-pathological contractions (Fig.5G). Additionally, GD/BUD restores propionate metabolism, which is linked to fatty acid synthesis from microbiota fermentation, essential for maintaining epithelial cell-cell and cell matrix adhesion and inhibiting epithelial to mesenchymal transitions associated with fibrotic states ^27,28^. Together, these molecular and pathway-level changes suggest that the combined inhibition of epithelial damage and inflammation by GD and BUD synergistically restores diseased airways toward a differentiated, more functional state.

## Discussion

We present a strategy to reverse airway remodelling in asthma by targeting the mechanical feed-forward cycle of epithelial damage. Rather than arising from ASM amplification and inflammation alone, hyperresponsiveness reflects a structural shift in smooth muscle architecture — from sparse, cross-hatched fibres to densely aligned parallel bundles reinforced by matrix, consistent with architecture observed in asthmatic macaque monkeys. This remodelling is mechanically driven: HDM triggers cyclical ASM contraction that pulls fibres into parallel bundles, enabling tighter constriction, while bronchoconstriction-induced epithelial damage fixes this configuration in place. We previously showed that bronchoconstriction causes excessive epithelial extrusion and damage, preventable with gadolinium and other extrusion inhibitors ^1^. The fact that gadolinium reverses established remodelling suggests that ongoing epithelial wounding locks ASM into a hyperresponsive state, possibly by signalling underlying smooth muscle to remain contracted as fewer epithelial cells cover the airway surface, analogous to stromal contraction in wound healing, and consistent with the prodromal airway tightening reported before asthma attacks ^29^. Together, this creates a feed-forward loop of epithelial damage and ASM remodelling, which gadolinium can block (Fig. 5J). Although we demonstrate reversibility in young mice, the fact that many children outgrow asthma ^30^ suggests airway responsiveness can be reset; adult or severely fibrotic airways may prove more resistant and warrant dedicated investigation.

Since asthmatic patients experience both allergen sensitisation and bronchoconstriction-driven exacerbation continuously, gadolinium or a similar mechanosensitive channel blocker could interrupt the mechanical damage driving both processes, creating a permissive environment for repair. Budesonide alone fails to break this cycle because it prevents neither epithelial damage nor the inflammation that follows an attack. Our transcriptomic and proteomic data show, however, that budesonide synergises with gadolinium, likely by calming baseline inflammation to accelerate structural recovery. Combining extrusion inhibitors with bronchodilators and anti-inflammatory agents could therefore address symptoms while achieving the structural repair that current therapies cannot.

Gadolinium is not without caveats. Chelated gadolinium injected repeatedly as an MRI contrast agent can bioaccumulate in damaged kidneys and the brain, though without known associated pathologies ^31^. Inhaled gadolinium salt, likely more toxic than chelated forms, was nonetheless well tolerated in our mice in the short term and did not accumulate in lungs or systemically. This transient, reversible aspect of gadolinium is critical since long-term suppression of extrusion can promote cancer ^32^. Moreover, as gadolinium is not specific to only Piezo1, its ability to target TRP channels, which are often mutated in severe asthma, suggests broader therapeutic potential.

With the usual caveats about safety and clinical translation, our findings indicate that locally and transiently inhaled inhibitors of epithelial extrusion could offer a route not merely to symptom control but to halting — and potentially reversing — the disease itself. Moreover, the ongoing wound healing that sustains asthma may also drive progression to lung cancer and COPD ^33^, suggesting that therapies targeting the epithelial damage cycle could deliver benefits well beyond asthma management.

## ACKNOWLEDGEMENTS

We thank Michael Redd, David Jackson, Joanna Porter, and Christopher Brightling for helpful comments on our manuscript. We are grateful to Sejal Seglani and the vibrant asthma research community in London, who have provided invaluable insight into this disease. We thank King’s College London Mouse Facilities, and Queen Mary University of London, King’s College London, Imperial College London, and University of Southampton Pathology Cores. We acknowledge the Genomics STP, and particularly Deb Jackson, Ashley Fowler and Marg Crawford for their contributions to RNA library preparation and sequencing, and Tegan Gilmore and the bioinformatics STP for access and processing of transcriptome at the Francis Crick Institute. We also like to thank Steven Lynham at King’s College London for proteomics analysis and Alex Morell and Piotr Golda at the London Metallomics facility at KCL for gadolinium measurements.

## AUTHOR CONTRIBUTIONS

J. R., D.C.B., and T. R. designed experiments, interpreted, analyzed data, and wrote the manuscript. D.C.B., and T. R. conducted all the experiments. S.M. analysed the alignment of airway smooth muscle. R.M.N. analysed the transcriptomics and proteomics data. All authors edited the manuscript.

## DATA AND MATERIALS AVAILABILITY

All data are available in the main text or the supplementary materials.

## SUPPLEMENTARY MATERIALS

Figures S1 to S5

Movies S1-S10

## Supplemental Figure Legends

**Supplemental figure 1.**
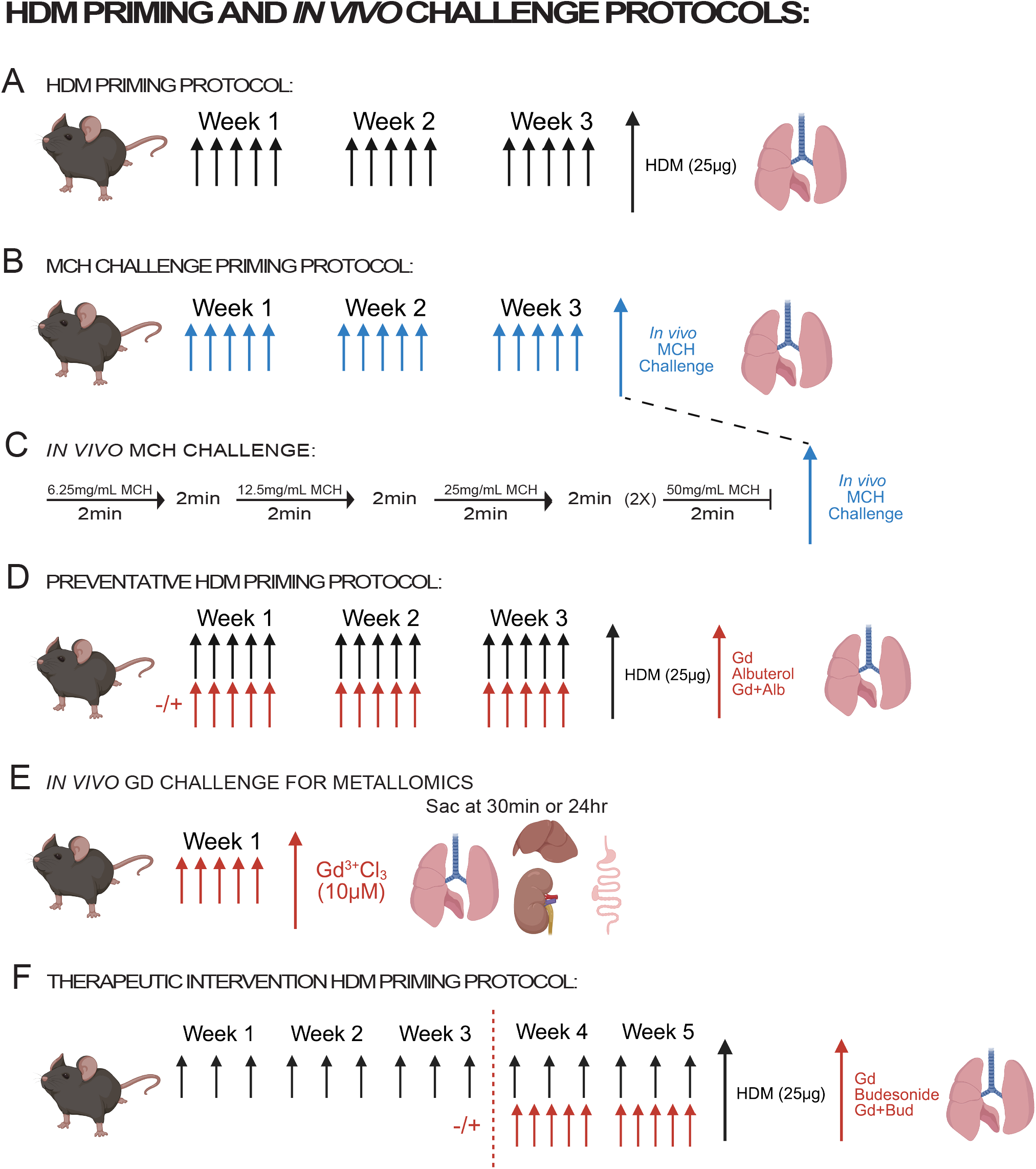
HDM priming and in vivo challenge protocols. (A) Priming schematic where mice received intranasal instillations of 25 µg HDM (black arrows) 5X per week for three weeks. (B) Bronchoconstriction-induced priming protocol where mice were challenged live with increasing dosages of MCH (blue arrows) 5X per week for three weeks. (C) Schematic of live MCH challenge used in (B) where dosages of MCH were doubled (from 6.25 to 50 mg/mL) for 2 min treatments, with 2 min rests in fresh air between challenges. (D) Preventative priming protocol where mice received HDM, as in (A), with or without GD (10 µM) and ALB (5mM) (red arrows). (E) In vivo gadolinium challenge where mice received consecutive 10 µM intranasal instillations of GD once per day for five days for metallomic analysis. (F) Schematic of therapeutic intervention HDM priming protocol where mice were exposed to HDM (black arrows) 3X per week for three weeks to establish asthma, before treating with GD, BUD, both (red arrows), or nothing for an additional two weeks along with HDM. Schematics were generated using BioRender.

**Supplemental figure 2.**
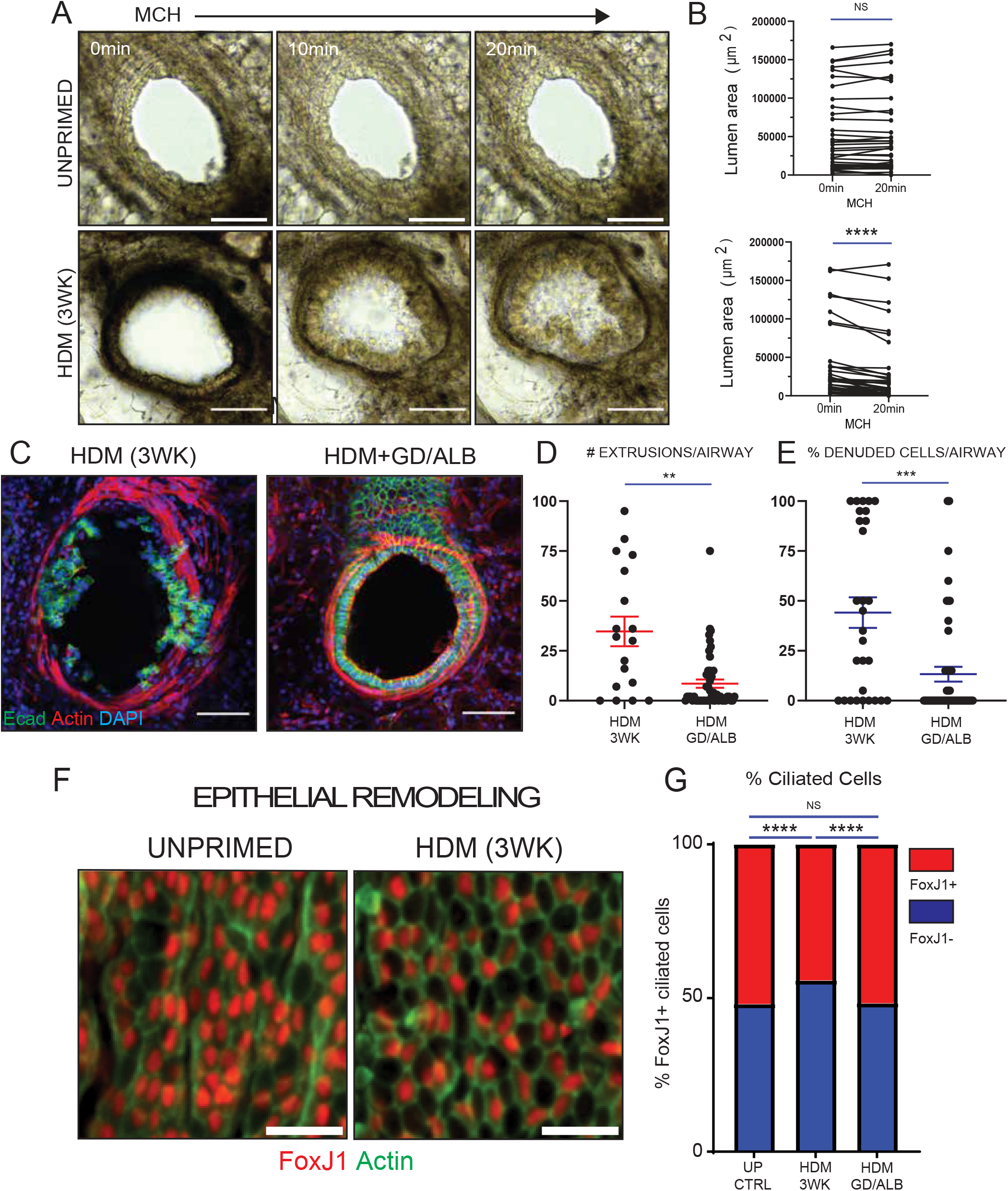
Gadolinium treatment at the onset of HDM priming prevent airway remodelling. (A) Movie stills from ex vivo lung slices from control and HDM primed mice treated with 500 mg/mL MCH (Scale bars, 50 µm), where constriction is quantified in (B) from measuring the lumen area at time 0 min and 20 mins following 500 mg/mL MCH treatment (unprimed: 43 airways from n=3; HDM (3wk): 40 airways from n=4, ****p<0.0001 from a Wilcoxon pairs-matched signed rank test). (C) Representative confocal projections of airways following ex vivo 500 mg/mL MCH treatment from mice exposed to HDM or HDM+GD/ALB for three weeks, immune-stained for the epithelium (Ecad), actin (phalloidin), and DNA (DAPI) (Scale bars, 50 µm). Quantification of epithelial cell extrusion (D) and precent denuding (E) per airway: HDM (3wk): 29 airways from n=3; HDM+GD/ALB: 48 airways from n=3 (**p<0.001, ***p<0.0005 from a Mann-Whitney test). (F) Confocal projections of ex vivo lung slice monolayers stained for FoxJ1 (ciliated cells) and actin, quantified in (G) for FoxJ1-positive cells (****p<0.0001 from a Chi-squared test). (H) Haematoxylin and eosin (H&E) stained lung sections from mice exposed to HDM or HDM+GD/ALB for 3 weeks (Scale bars, 500 µm), scored for inflammation degree per airway in (I), where 0 is little to no immune cell infiltrate and 3 is severe inflammation (n=6 per treatment group; *p<0.05 from a Chi-squared test).

**Supplemental figure 3.**
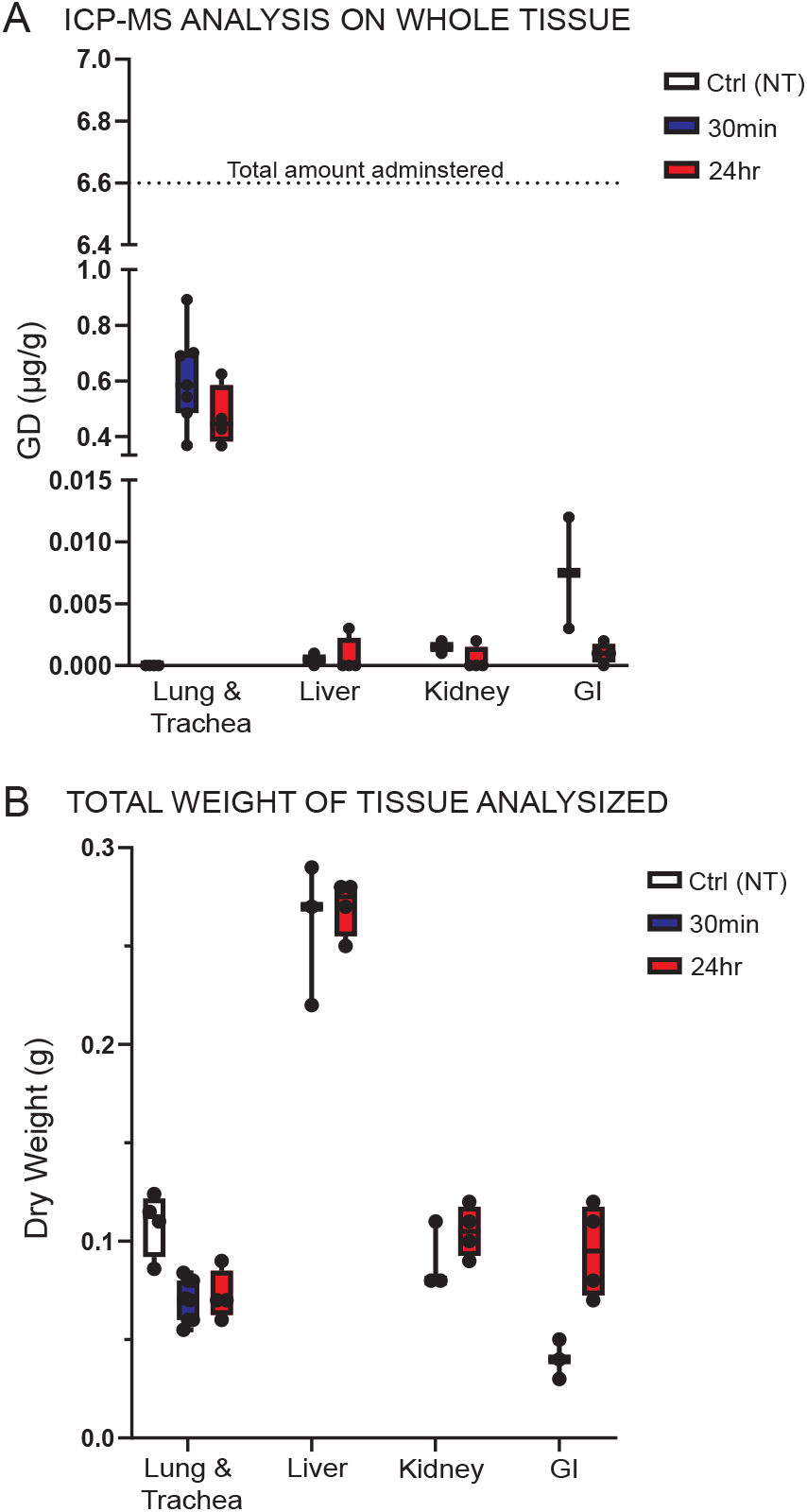
Metallomics analysis showing repeated doses of non-chelated gadolinium do not accumulate in the lung or other tissues. (A) ICP-MS analysis of total gadolinium concentrations from tissue isolated from mice intranasally treated with 10 uM gadolinium (3+) hexa-chloride as described in fig. S1E (control: lung, n=4), which were measured at 30 min (lung: n=8; liver: n=3; kidney: n=3; GI: n=3) and 24 hrs (lung: n=4; liver: n=4; kidney: n=4; GI: n=4) after the finial instillation. (B) Average weight of total dry tissue used for metallomic measurements.

**Supplemental figure 4.**
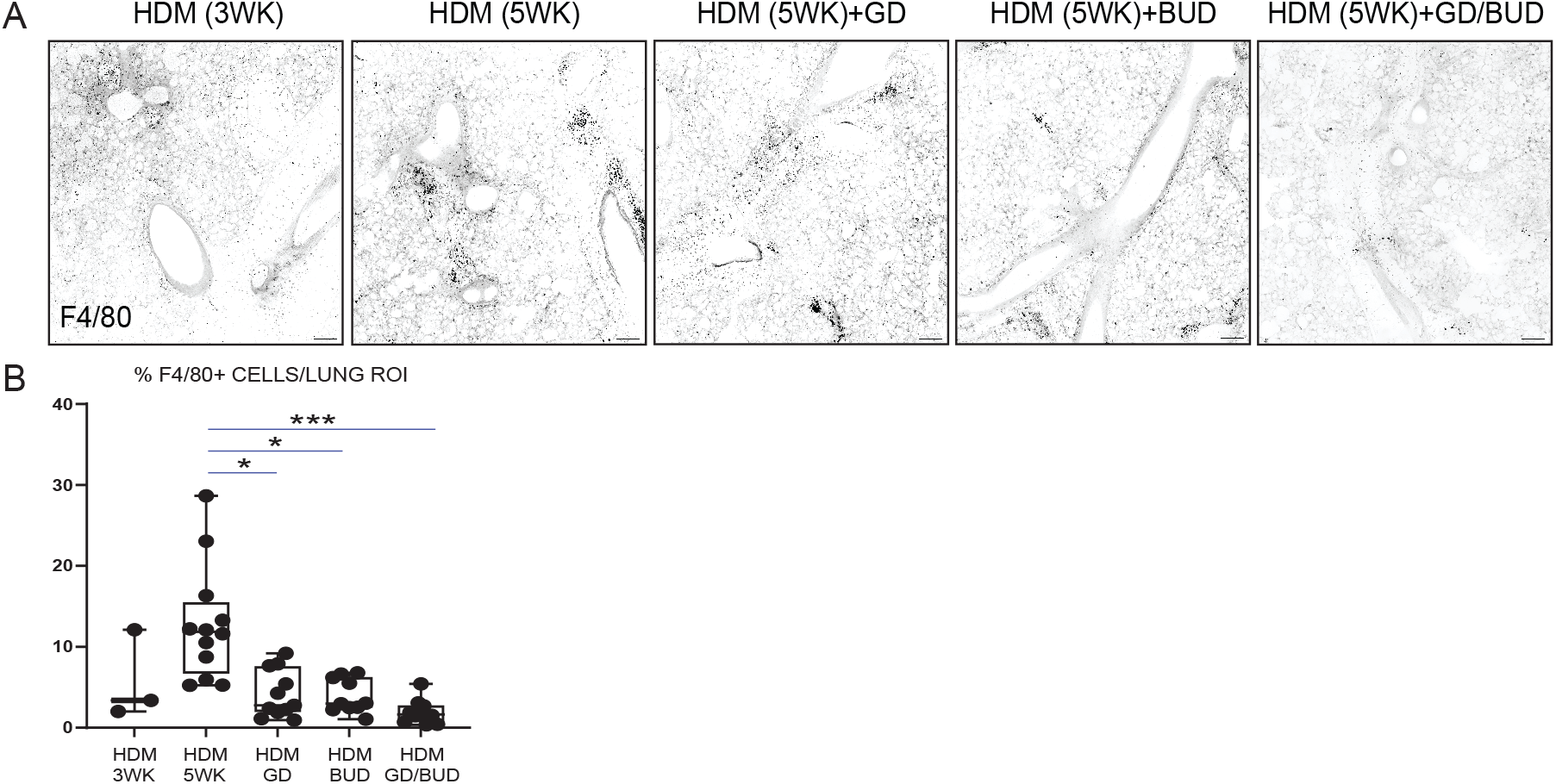
Gadolinium and budesonide cooperate to reduce the number of macrophages in asthmatic lungs. (A) Representative large confocal projections from 16 20X fields stitched together (with 10% overlap between images) of immune-stained lung slices for macrophages (F4/80) from mice primed with HDM mice ± treatments, quantified in (B) as percentages of F4/80-positive cells in lung slices (n=3 for all groups analysed, *p<0.05, ***p<0.0005 from a Kruskal-Wallis test test) (Scale bars, 200 um).

**Supplemental figure 5.**
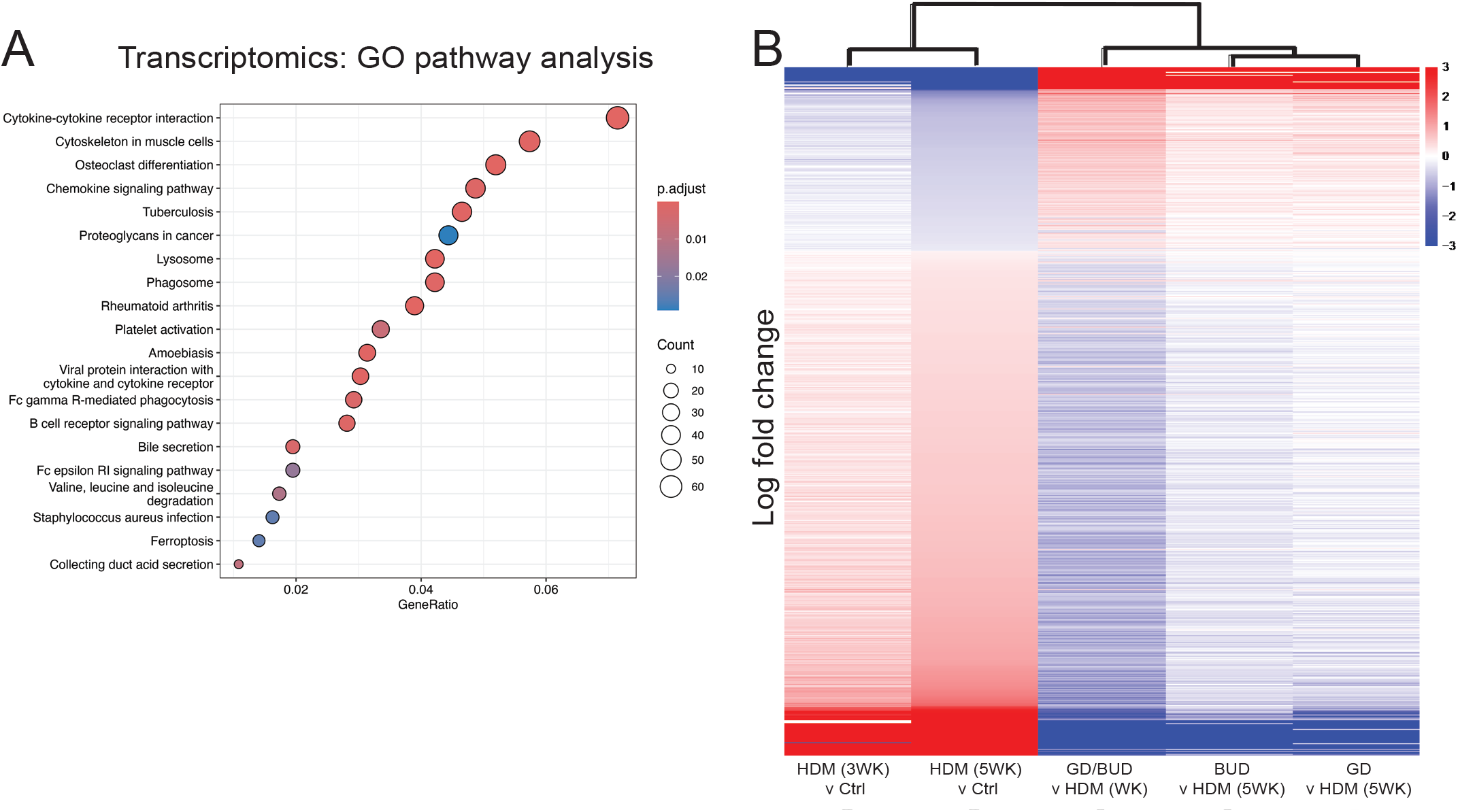
Gadolinium and budesonide work synergistically to reverse asthmatic molecular landscapes. (A) Dot plot showing the enrichment in KEGG pathways that the differentially expressed genes in A mapped to, depicting p-adjusted values (red-blue scale) and number of genes (dot size) per pathway. (B) Hierarchical heatmap (log2 fold changes) of the 730 proteins intersecting between 5-weeks of HDM priming and gadolinium + budesonide treatment (HDM5 vs control, p-adjusted < 0.05) showing reversed differential protein expression with HDM5 versus gadolinium + budesonide treatment.

**Supplemental figure 6.**
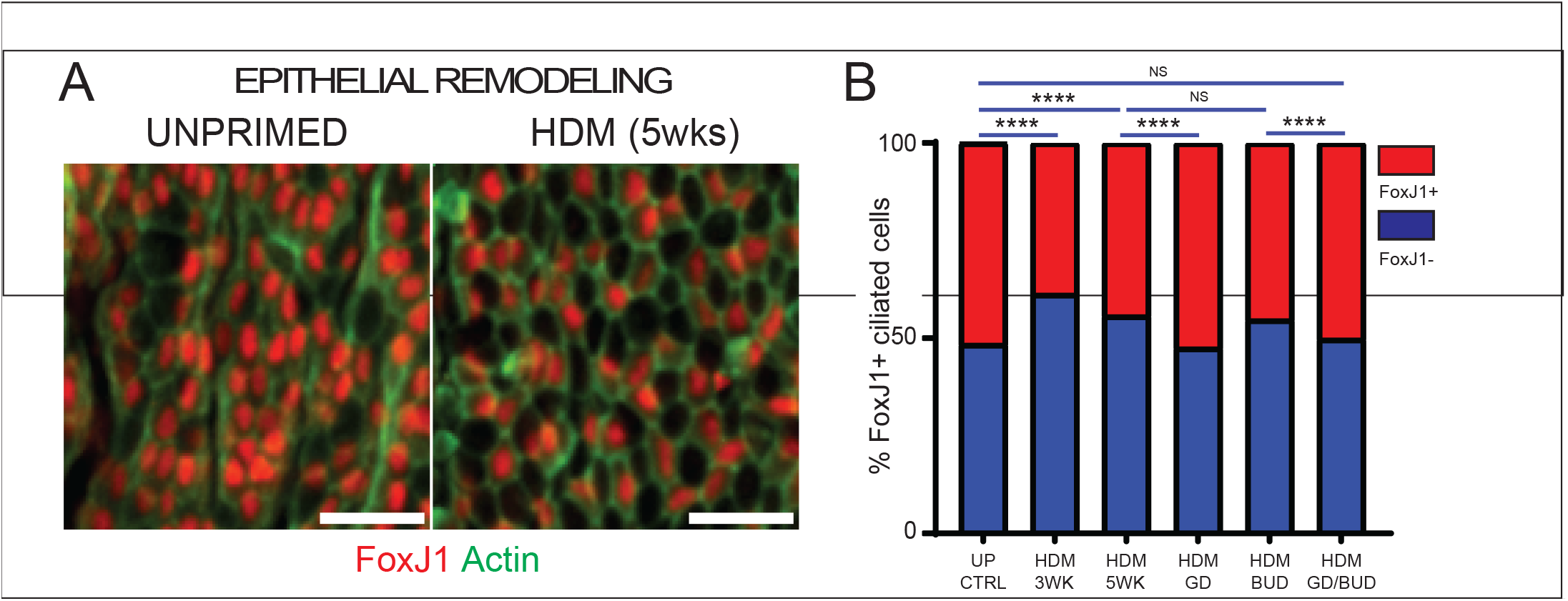
Gadolinium reverses loss of FoxJ1+ ciliated cells. Epithelial cell types restored by GD/BUD priming. Representative ex vivo monolayers immune-stained for FoxJ1 and actin to determine the number of ciliated (FoxJ1+) cells from each treatment group (n=3; ****p<0.0001 from a Chi-squared test) (Scale bars, 25 um).

## Supplemental Movie Legends

**Supplemental movie 1. Unprimed healthy ex vivo lung slices respond mildly to MCH treatment**. Bright field live imaging of a lung slice from an unprimed control mouse treated with 500 mg/ml MCH for 20 mins at 30 sec intervals (Scale bars, 50 μm).

**Supplemental movie 2. Live imaging of healthy lung slices reveal a robust mechanical response to HDM exposure**. Live imaging of 1μg/μL HDM treatment in a small unprimed airway for 60 mins at 15 sec intervals (Scale bars, 50 μm).

**Supplemental movie 3. Live imaging of healthy ex vivo lung slices reveal a robust mechanical response to HDM exposure in all airway sizes**. Representative movie of healthy airways of all sizes responding to 1μg/μL HDM treatment for 30 mins at 30 sec intervals (Scale bars, 50 μm).

**Supplemental movie 4. Healthy lung slices respond mildly to MCH treatment**. Live imaging of a lung slice from an unprimed control mouse treated with 500 mg/ml MCH for 20 mins at 30 sec intervals (Scale bars, 50 μm).

**Supplemental movie 5. HDM primed lung slices hyperconstrict to MCH treatment**. Movie of a representative airway from a mouse primed with HDM for three weeks (fig. S1A) treated with 500 mg/mL MCH for 20 mins at 30 sec intervals (Scale bars, 50 μm).

**Supplemental movie 6. Gadolinium, with albuterol, treatment protects against MCH-induced epithelial damage**. Representative movie of an airway treated with 500 mg/mL MCH for 20 mins at 30 sec intervals from a mouse primed with HDM+GD/ALB for three weeks (fig. S1D).

**Supplemental movie 7. Airways primed with HDM for 5 weeks are hyperresponsive**. Live imaging of an airway from a mouse primed with HDM for 5 weeks challenged with 500 mg/mL MCH for 20 mins at 30 sec intervals.

**Supplemental movie 8. Gadolinium treatment reverses airway hyperresponsiveness in HDM primed lung slices**. Live imaging of an airway from a mouse primed with HDM for 3 weeks before gadolinium with HDM for another 2 weeks challenged with 500 mg/mL MCH for 20 mins at 30 sec intervals.

**Supplemental movie 9. Budesonide does not prevent hyperresponsiveness during HDM priming**. Live imaging of an airway from a mouse primed with HDM for 3 weeks before budesonide with HDM for another 2 weeks challenged with 500 mg/mL MCH for 20 mins at 30 sec intervals.

**Supplemental movie 10. Gadolinium, with or without corticosteroid, treatment reverses airway hyperresponsiveness in HDM primed lung slices**. Live imaging of an airway from a mouse primed with HDM for 3 weeks before gadolinium and budesonide with HDM for another 2 weeks challenged with 500 mg/mL MCH for 20 mins at 30 sec intervals.

## Materials & Methods Animal Model

House dust mite (HDM) priming: 6-to 8-week-old C57BL/6J mice (C57BL/6JOlaHsd from Envigo, UK) were briefly anaesthetised with isoflurane and had 25µg (total protein) of HDM (D. pteronyssinus – Citeq Biologics, Groningen, The Netherlands) dissolved in 25µl of PBS intranasally instilled for either 3 or 5 weeks, with or without therapeutic interventions at 3 weeks, according to the priming protocol (fig. S1). Mice were sacrificed 24 hours after the final challenge by rising concentration of CO_2_, confirmed by exsanguination of the femoral artery. Mice numbers are listed in figure legends of appropriate experiments and were determined using NC3R’s principles.

All animals were maintained under specific pathogen–free conditions and handled in accordance with the Institutional Committees on Animal Welfare of the UK Home Office Animals (Scientific Procedures) Act 1986. All animal experiments were approved by the Ethical Review Process Committee at King’s College London and carried out under license from the Home Office, UK. All mice were humanely handled – Program Project License P68983265 & PP8646082 to Jody Rosenblatt at King’s College London.

### In vivo methacholine (MCH) Challenge

Mice either received this challenge 5 times per week for 3 weeks or once at the end of their immune-priming protocol. Mice were placed into a 6.5-quart Hefty bin fitted with an Aerogen Pro Nebulised System (AG-AP6000-XX) that produces 2.5-4.0µm volume mean diameter aerosolised particles ^1^. Mice were nebulised with two-minute challenges of increasing concentrations of MCH (6.25, 12.5, 25 and finally 2 × 50 mg/mL), resting in fresh air for 2-minutes between the challenges. Mice were sacrificed either 30 minutes or 24 hours after the final MCH challenge by rising concentration of CO_2_, confirmed by exsanguination of the femoral artery.

### Precision-cut ex vivo Lung Slices (PCLS)

Ex vivo lung slices were obtained from mice within 48 hours of their last challenge. The chest cavity was opened and the trachea carefully exposed, where a small incision was made to accommodate the insertion of a 20Gx 1.25 canula (SURFLO I.V. catheter). The lungs were inflated with 2% low melt agarose (Fisher – BP1360) dissolved in HBSS+ (Gibco – 14025), after which the lungs, heart and trachea were excised. Washed in sterile PBS, and lobes separated into their individual components. Individual lobes were embedded in 4% low melt agarose and solidified on ice for 15 minutes. A Leica VT1200S vibratome was used to cut 200-micron thick slices, which were washed and incubated in DMEM/F-12 medium supplemented with 10% fetal bovine serum (FBS) and 1% penicillin/streptomycin. The ex vivo lung slices remained viable (MCH-reactive) for at least one week after isolation ^1,34^.

### Ex vivo house dust mite (HDM) and methacholine (MCH) treatment with live imaging

Ex vivo lung slices were experimentally treated with 1 μg/μL HDM or 500 mg/mL MCH (acetyl-**β**-methylcholine chloride, Sigma A2251) for 20 or more mins in Hank’s buffered saline solution (HBSS+). MCH is a non-selective muscarinic receptor agonist clinically used to test the severity of asthma in patients. Lungs were incubated in HBSS+ in 24-well plates at 37°C with HDM or MCH for 20+ minutes and filmed at 30 sec intervals using a Life Technologies EVOS FL Auto microscope to measure bronchoconstriction responses. For airway constriction measurements in responses to treatment, the lumen area (the “dead” or empty space in the airway lumen) at 30 sec intervals (HDM), or at time=0 min and at time=20 min (MCH) was measured using Fiji software from live imaging stills.

### Immunofluorescence and imaging of fixed PCLS

PFA-fixed ex vivo lung slices were incubated for one hour at room temperature in blocking solution: PBS containing 0.1% triton X-100, 0.1% sodium azide, and 2% bovine albumin (BSA), before incubating overnight at 4°C at 1:100 in blocking solution for all primary antibodies used: rabbit anti-E-Cadherin (24E10 – Cell Signaling); rabbit anit-FoxJ1 (EPR21874 - Abcam); mouse anti-Tubulin, acetylated (T6793 – Thermo Scientific); rat anti-F4/80 (CI:A3-1 – Bio-Rad). Ex vivo lung slices were washed 3 × 30 minutes in PBS+0.5% Triton X100) before incubating overnight - A11008) or anti-mouse (A32723) IgG, Alexa Fluor 568 goat anti-rabbit (A11011) or anti-mouse (A11004) IgG, or Alexa Fluor 647 goat anti-rabbit (A32733) or anti-mouse (A21235) IgG, or Alexa Fluor 647 goat anti-rat (AB150159 – Abcam) + 1:250 Alexa Fluor 488, 568, or 647 Phalloidin (Thermo Scientific – A12379, A12380, A22287, respectively). Slices were washed 2 × 30 minutes in PBS+0.5% Triton X-100, stained with 1:1000 DAPI in PBS for 20 minutes, mounted in ProLong Gold (Invitrogen P36930), and imaged on a Nikon Eclipse Ti2 spinning disc confocal microscope with a 20X or 40X objective.

### Airway smooth muscle alignment, cross-hatching and thickness quantifications

Using a methodology described by Marcotti et al. ^35^, Alignment by Fourier Transform (AFT), images of ASM banding around the airway were analysed using MATLAB (Mathworks, v2023b) to quantify feature alignment. The script is open-source and available at https://github.com/OakesLab/AFT-Alignment_by_Fourier_Transform. Maximum intensity projections of images of ASM were converted into 8-bit format. A user-drawn binary mask was applied to the images to restrict the analysis to the airway only. AFT parameters were set as follows. An intensity threshold was set to 20 (0-255) to exclude background noise. The analysis windows were set to a size of 75 pixel with a 75% overlay. Alignment was investigated over a radius of two neighbourhoods around each reference window. The median alignment score across the whole airway was reported, with 0 representing random alignment and 1 perfect alignment of the ASM bundles. Percentage cross-hatching of ASM bundles was calculated by counting the number of ASM bundles and the number of ASM bundles which intersect with another ASM bundle across a whole airway. ASM augmentation was calculated by averaging the distance between actin bundles per airway.

### Histological analysis and scoring

At the end of their treatments, mice were humanely euthanized by CO_2_ inhalation followed by cervical dislocation or exsanguination via the femoral artery. The lungs were inflated with 10% neutral buffered formalin (NBF), excised, and fixed in NBF overnight at room temperature, followed by another overnight room temperature incubation in 70% ethanol. The large left lung was excised and embedded in paraffin. Using a keratome, 3 × 5 µm thick serial slices were made per slide, 3 slides per lung, each slice made at 50 µm deeper intervals and stained with haematoxylin and eosin (H&E) or Mason’s trichrome. Slides were scanned at 20X magnification using a Zeiss AxioScan.Z1. Inflammatory scoring was done as previously described ^1^, where each airway received a score of (0) for little-to-no immune infiltrate up to (3) for severe inflammation. Basement membrane and ASM thickness was determined by the average of 4 measurements per airway.

### ICP-MS Metalomics Analysis

Whole tissue was harvested and stored at -80°C until analysis. Tissue samples were transferred to pre-weighed trace grade centrifuge tubes (ElkayLabs) and allowed to dry overnight in an oven at 70°C. Samples were cooled and weighed before being digested by the addition of 0.7 mL Optima grade concentrated nitric acid (67–69% wt/wt; Fisher Scientific) and 0.1 mL hydrogen peroxide (30% w/w, Supelco) followed by heating at 60 °C in a heat block for 1 h. Digested samples were diluted up to 14 mL using purified water with a resistivity of ≥18.2 MΩ cm (Milli-Q IQ 7015, Merck, Massachusetts, USA) and spiked with Ga (100 mg/L) as the internal standard to achieve a final concentration of 10 µg/L.

All measurements were conducted on a Perkin Elmer NexION 5000 Inductively Coupled Plasma Triple Quadrupole Mass Spectrometer under Dynamic Reaction Cell (DRC) mode at the London Metallomics Facility, King’s College London. The introduction system to the instrument was a Cetac ASX-560 autosampler coupled to a SeaSpray glass nebulizer fitted to a quartz cyclonic spray chamber. RF power was 1600 W, Argon plasma flow and nebulizer gas flow rates were 18 L min-1 and 0.98 L min-1, respectively. A calibration curve was prepared by serial dilutions of an ICP-MS Multi-element standard (Merck) in a range of 0.10 – 500 ppb of Calcium (^40^Ca) measurements were taken in DRC mode with a 0.6 mL/min flow of Ammonia inside the reaction cell and quadrupole RPq of 0.7 and Gadolinium (Mass shift ^157^Gd – ^157^Gd^16^O) taken in mass shift DRC mode with a 1 ml/min flow of oxygen inside the reaction cell and quadrupole RPq of 0.7. Integration time for the analyte’s was 1000 ms and samples were measured in 5 instrument replicates.

Quality control of ICP-MS measurements was ensured through a combination of repeat measurements of acid blanks, a calibration standard, and a certified reference material (CRM) repeated at intervals during the analysis.

Analyte measurements were normalized to the internal standard ^71^Ga to account for instrument drift and matrix effects, and measurements were subsequently blank corrected by removing the average analyte intensity of repeat blank measurements. The corrected isotopic intensity was converted to concentration measurements by regression analysis using the calibration curve. The quality of the regression analysis was confirmed by verifying the linearity of the calibration curve.

### Transcriptomics

#### Sample preparation

The large left lobe was excised and snap frozen in liquid nitrogen in Trizol and stored at -80°C until RNA could be isolated. Lung tissue was homogenized using a MP Biomedicals FastPrep-24 5g bead beating grinder lysis Sample Prep Homogenizer using 3 rounds of 6m/sec beating for 40 secs with 2 min incubations on ice between cycles, using 3 mm tungsten carbide beads (Qiagen – 69997). Homogenate was passed through a Qiagen Shredder column before using a RNeasy Plus Mini kit (Qiagen – 74134) with DNase treatment per manufacturer’s instructions to isolate high quality RNA.

#### Library preparation and sequencing

Watchmaker Genomics RNA libraries were prepared manually following the manufacturer’s protocol (Combined Protocol: Watchmaker RNA Library Prep Kit with Polaris™ Depletion, version v1.1.1122) for the Watch-maker RNA Library Prep Kit (7K0078-096) and Polaris Depletion Kit - rRNA/Globin (HMR, 7K0077-096).

Samples were normalized to 100ng total RNA material per library in 18μl of nuclease-free water. 4.5µl of Depletion Master Mix and 2.5µl Depletion probes – rRNA/Globin (HMR) were added to each sample and incubated at 77oC for 2 minutes then at 65oC for 15 mins. Bound probes were digested by addition of 35µl Probe Digestion Master Mix, followed by incubation at 37oC for 10 minutes. Depleted samples were cleaned up with SPRISelect beads (ratio: 1.8x).

Samples were combined with 27µl Frag and Prime Buffer and incubated at 85oC for 5mins to fragment the RNA to approximately 200nt.

To perform first strand synthesis, 25µl of fragmented RNA was combined with 9µl of First Strand Buffer and 1µl of First Strand Enzyme. Samples were incubated at 25oC for 10mins followed by 15mins at 42oC, then 15mins at 70oC to inactivate the reaction. To perform second strand synthesis and A-Tailing, 14µl of Second Strand Buffer and 1µl Second Strand Enzyme were added to each sample and incubated at 42oC for 5 minutes, followed by 62oC for 10 minutes. Adaptors were ligated to the cDNA fragments by adding 40µl Ligation Buffer, 5µl Ligation Enzyme and 5µl Adaptor (diluted to 4µM), then incubating at 20°C for 15 minutes. Adaptor-ligated samples were cleaned up with SPRISelect beads (ratio: 0.7x). For the amplification of the library, 25µl of Equinox Amplification Master Mix was added, plus 5µl of a unique index (xGen UDI 10nt Primers [IDT 10008052]). The number of PCR cycles applied for library construction were 7 according to the manufacturer recommendation for 100 ng input DNA. Final libraries were cleaned up with SPRISelect beads (ratio: 1x). The quality of the purified libraries was assessed using an Agilent D1000 ScreenTape Kit on an Agilent 4200 TapeStation. Libraries were sequenced to a depth of at least 24M reads on an Illumina NovaSeq X run in 101-8-8-101 configuration.

#### RNAseq data analysis

RNA was preserved in TRIzol and extracted according to manufacturer’s instructions, with DNAse treatment prior to library with polyA selection and sequencing. FASTQ files were quality checked and mapped against the M.musculus transcriptome (GRCm39). Samples were tested for batch effects. To compare all 6 groups of mice we performed DESeq2 analysis using the likelihood ratio test (LRT) model, better suited to our experimental design that included 6 groups. We compared all groups to house dust mite treated animals for 5 weeks, and house dust mite treated mice for 5 weeks to control mice. As this model is less restrictive statistically than the standard DESeq2 pipeline using Wald tests, we restricted our analysis to p-adjusted values of less than 0.001, resulting in 1940 genes differentially expressed.

### Proteomics

#### Sample preparation

Whole mouse lungs were excised, placed into a 1.5ml Eppendorf and flash frozen in liquid nitrogen before being transferred to a -80°C freezer for sample preparation. Each sample was transferred to 2ml Lo-bind centrifuge tubes and washed with 200ml of 0.2M EPPS pH8.5 (Cat # J61476.AK; Thermo Scientific) and briefly vortexed to remove residual blood from the tissue. This was repeated for a total of three times with supernatant removed to waste each time. The EPPS was removed to waste and replaced with 1ml of 8M Urea with protease (Complete Mini; #11836170001; Merck) and phosphatase (PhosStop, #4906845001; Roche) inhibitors included.

Samples were subjected to sonication using an ultrasonic probe set at 3 watts for 3 second bursts in four cycles to prevent overheating (Branson Sonifier 150). Samples were always kept on wet ice during the process. Following sonication, samples were centrifuged at 14,000 rpm for 2 minutes to pellet non-solubilised material. The pellet was drawn five times through a 23G gauge needle and the solution vortexed well. The sample was centrifuged at 14,000rpm for 2 minutes to check for residual pelleted material. Once solubilisation was complete, protein concentration was determined using a Nanodrop One system (Thermo Fisher Scientific) set at A280 protein with 340nm baseline correction and mg/ml units. The full measurements for each sample are represented in table S1 in Appendix 1. All samples were normalised to 100mg total protein in 100ml of 8M urea in new Lo-Bind 1.5ml centrifuge tubes for further processing.

#### Reduction and alkylation

Samples were reduced using tris(2-carboxyethyl)phosphine hydrochloride, TCEP (Sigma-Aldrich C4706, MW 286.65) at a concentration of 50mM in 0.2M EPPS pH8.5. Added 11ml of 50mM DTT to give a final concentration of 5mM DTT, vortex then centrifuge briefly at 14,000 rpm. Incubate at room temperature, room temperature (RT) for 30 minutes.

Following reduction, samples were alkylated with iodoacetamide, IAM (Sigma-Aldrich A3221, MW 184.96) at a concentration of 20mM in 0.2M EPPS taken from a stock solution of 200mM in 0.2M EPPS. Added 12ml of 200mM IAM to give a final concentration of 20mM in a 4:1 ratio of excess to TCEP. Incubate at RT in the dark for 30 minutes.

To prevent unwanted alkylation reactions with amino acids other than cysteine, the IAM was quenched briefly (10-minute incubation at RT) with 11ml 50mM dithiothreitol, DTT (Sigma-Aldrich D5545, MW 154.25) in 0.2M EPPS pH8.5. Samples were vortexed and centrifuged.

Proteins were precipitated to remove the lysis buffer and excess reduction & alkylation agents. To each tube, added 400ml ice-cold methanol, then added 100ml ice-cold chloroform and finally 300ml ddH_2_O. Tubes were vortexed well then centrifuged at 14,000 rpm for 1 minute to produce a protein disk between the aqueous and solvent layers. The liquid was carefully removed to avoid disruption to the protein disk. Added 400ml ice-cold methanol to wash the sample prior to centrifugation at 14,000 rpm for 1 minute. Remove the methanol. The protein disk was resuspended in 100ml 0.2M EPPS pH8.5.

Enzymatic digestion was performed using trypsin (Roche, Cat# 11418475001) in 0.2M EPPS to cleave at arginine and lysine residues (except where the residues lie adjacent to a proline) in a 1:100 enzyme:substrate ratio (1mg total enzyme). Samples were incubated at 37°C overnight.

25mg of each sample was removed to new Lo-bind tubes and dried to completion in s Speed Vac (Thermo Fisher) and resuspended in 50ml of resuspension buffer (2% acetonitrile (ACN) in 0.05% trifluoroacetic acid (TFA)) for a final concentration of 0.5mg/ml.

#### LC-MS/MS

Samples were injected as 2ml (1mg concentration on column) for chromatographic separation using a U3000 UHPLC NanoLC system (Thermo Scientific, UK). Peptides were resolved by reversed phase chromatography on a 75μm C18 column (50cm length) using a three-step linear gradient of 80% acetonitrile in 0.1% formic acid. The gradient was delivered to elute the peptides at a flow rate of 250nl/min over 60 min. The eluate was ionised by electrospray ionisation using an Orbitrap Eclipse (Thermo Scientific, UK) operating under Xcalibur v4.7.69.37. The instrument was first programmed to acquire data-independent acquisition (DIA) using an Orbitrap-Orbitrap Velocity method under higher-energy collision induced dissociation (HCD) fragmentation defined by short m/z windows.

#### Database Searching

Raw mass spectrometry data were processed DIA-NN software ^36^ using a library-free search against the Uniprot Mouse database (17,963 entries; March 2024), allowing for static carbamidomethyl (C) and variable oxidation (M) with stringency set at 1% FDR across the sample set.

Proteomics analysis was performed using limma, adding one count to enable analysis of proteins that appeared or disappeared between groups. Multiple test correction and P adjustment was performed in all cases. All p values noted in the figures correspond to p adjusted metrics. Plots were generated in R and pathway analysis was done employing KEGGREST and plotting using KEGGgraph.

## Data Availability

The mass spectrometry proteomics data have been deposited to the ProteomeXchange Consortium via the PRIDE ^37^ partner repository with the dataset identifier PXD068181.

